# Genome Sequencing, Molecular Marker Development and Genetic Diversity Assessment of Economically Important Vulnerable Tree Species *Saraca as*oca (Roxb.) W.J de Wilde

**DOI:** 10.1101/2025.01.17.633512

**Authors:** P Mahalakshmi, Santhosh N Hegde, Malali Gowda, Abdul Kareem, M Anbarashan, S Noorunnisa Begum, Subrahmanya Kumar Kukkupuni, Pavithra Narendran

## Abstract

*Saraca asoca*, is an understory tree along streams in evergreen to semi-evergreen forests up to 600 m. It is an important tree in cultural tradition and medicinally significant. It is native to India and Sri Lanka. Globally the species is found to occur in India, Sri Lanka, Myanmar, Bangladesh. It was introduced in Malaysia. Within India, it’s found in Western Ghat and Eastern Ghat. It is occasionally planted in gardens. *Saraca asoca* is known for its extensive pharmacological properties, particularly its bark is used in treating menorrhagia, dysfunctional uterine bleeding, hemorrhagic dysentery, and other gynecological issues, it holds a prominent place in Ayurvedic medicine. This study was undertaken to sequence the whole genome of *Saraca asoca* using the Illumina HiSeq2500 platform, and evaluating the genetic diversity among samples from Kolluru and other locations in southern India through Genotyping by Sequencing (GBS). Analysis of 49 samples established genetic diversity relationships using a distance matrix. Sequencing yielded 1.6 Gb, covering 76% of the estimated genome size. The genome includes 764 million bases of repetitive DNA elements. A survey of Simple Sequence Repeats (SSRs) identified 584,615 SSRs, with 236,123 sequences containing SSRs. Utilizing the KEGG database, biosynthesis pathways for catechin and epicatechin within the flavonoid synthesis pathway were identified. This comprehensive genomic analysis of *Saraca asoca* (Sita Ashoka) provides critical insights for conservation efforts aimed at preserving this vulnerable species, *Saraca asoca* (Roxb.) Willd.

## 1. Background

*Saraca aso*ca (Roxb.) Wilde (hereafter called as *Saraca asoca*), is one of the ancient tree species identified in India, with its medicinal heritage dating back to 5000 BC from one of the two important Indian epics Ramayana, thus commonly known as “Sita Ashok” also marking the significance of the birth of Buddha under the Saraca asoca tree (Bhalerao et al. 2014), depicting its widespread cultural heritage across north-east India and other parts of the world. The wild presence of this species has been recorded only from a few scattered patches in the Western Ghats of Maharashtra, Goa, Karnataka, Tamil Nadu, Kerala, and the Eastern Ghats of Odisha and Meghalaya (Yadav and Sardesai 2002, Sasidharan 2004, Singh and Karthikeyan 2000, Ved at al. 2016) **(Figure 1)**, though originated from India also found in Sri-Lanka, Bangladesh and other neighboring countries, at an altitude of up to 750m (Ramesh et al. 2016). Various parts of *Saraca asoca* **(Figure 1)** are identified for treating different types of ailments, in particular the bark of *Saraca asoca* is known to contain glycosides, flavonoids, tannins, saponins, alkanes, esters and primary alcohols and is used to treat menorrhagia, leucorrhoea, dysfunctional uterine bleeding (Mohan et al. 2016, Singh et al. 2015). Attributing to its usage in almost all the ayurvedic formulations and pharmaceutical industries. Besides gynecological complications *Saraca asoca* is effective in treating bacterial infections, skin problems, tumors, worm infestations, cardiac and circulatory problems in general (Singh et al. 2015).

**Figure 1:**
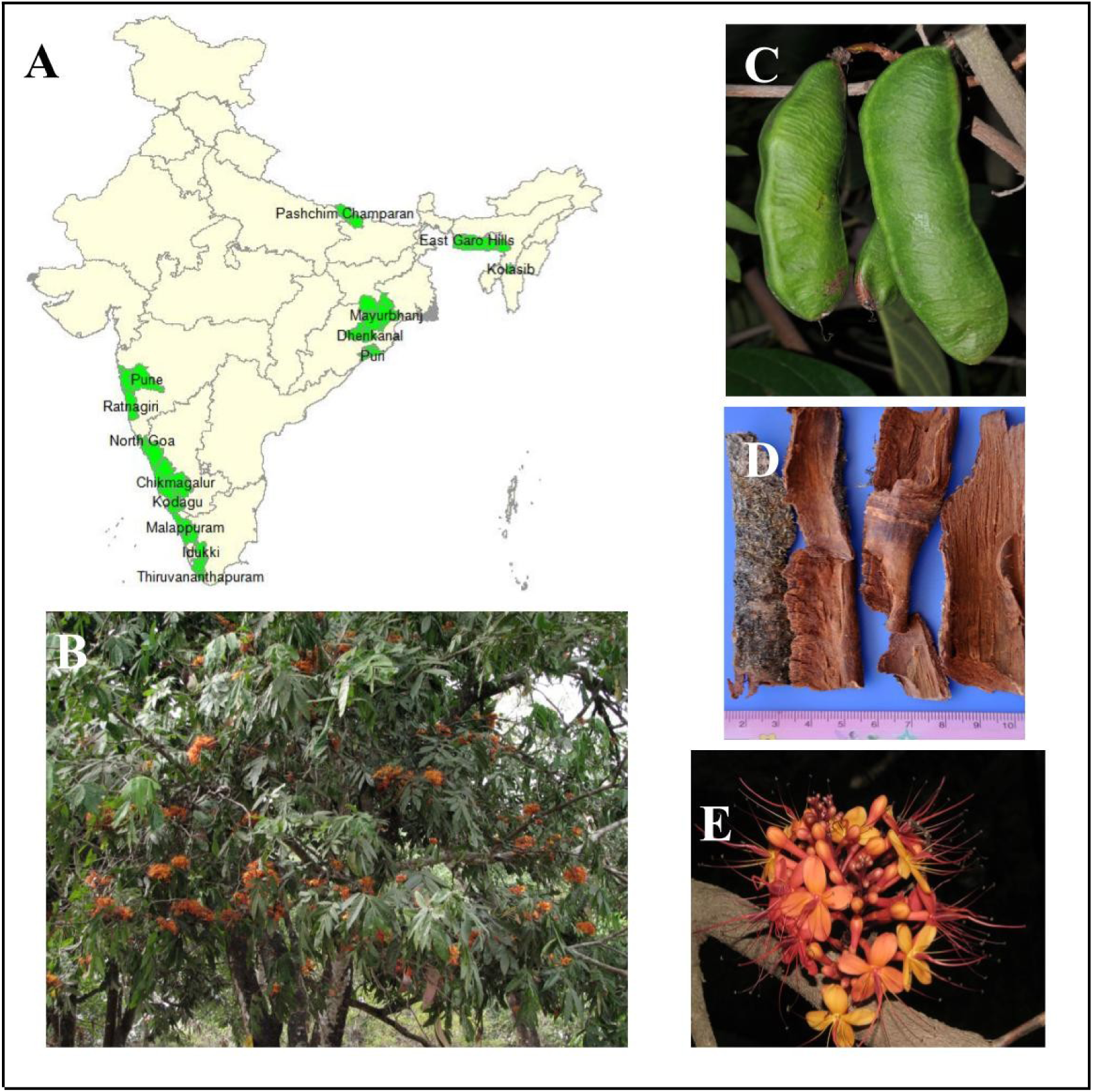
A – Distribution of *Saraca asoca* in India; green – distribution of *Saraca asoca* (source: https://envis.frlht.org/index.php/bot_search); B – *Saraca asoca* at The University of TransDisciplinary Health Sciences and Technology, Bengaluru, India; C – *Saraca asoca* fruit (green); D – *Saraca asoca* stem and bark; E – *Saraca asoca* flowers

The significant medicinal benefits of *Saraca asoca* have led to its premature felling by unethical sources, causing a sharp decline in the population of these trees. The rising demand for *Saraca asoca* bark has also led to an increase in product adulteration with this, however, Reportedly substituted/adulterated with plant materials obtained from Humboldtia vahliana Wight (Caesalpiniaceae: N.Sashidharan, KFRI, pers. commun.), Shorea robusta Gaertn. (Dipterocarpaceae) (Vaidya 2005) and Mallotus nudiflorus (L.) Kulju & Welzen (Euphorbiaceae) (Anon 2024). Consequently, *Saraca asoca* is classified as vulnerable by the International Union for Conservation of Nature (IUCN), (IUCN (2011) IUCN red list of threatened species. Version 2011.1, Smitha and Thondaiman, 2016). Therefore, the replenishment and conservation of *Saraca asoca* are of utmost priority. Hence this species has been conserved in situ in India as Medicinal Plant Conservation Areas (MPCA) by the forest departments.

The recent advances in genome sequencing through next generation sequencing (NGS) technologies are generating large number of sequences in affordable cost, thereby aiding in the development of molecular markers including SSRs particularly in non-model plants (Unamba et al. 2015), also providing opportunities to develop millions of novel markers, this approach also allows for the identification of different numbers of repeat motifs within SSRs (Panahi et al. 2024). The current study provides details of the molecular markers (SSR) from the whole genome of *Saraca asoca* for assessing genetic diversity and structure in *Saraca asoca* germplasm, and provides potentially useful information for developing conservation and breeding strategies for economically important vulnerable tree species *Saraca asoca*.

In the current research, the whole genome sequence data along with the biosynthesis pathway of catechin, epicatechin metabolites and the SSR marker study was carried out for the whole genome of *Saraca asoca.* Thus the present study could contribute to filling the research gap in the genomic understanding of *Saraca asoca,* an economically important vulnerable tree species. Also, with the next-generation sequencing (NGS) technology, we are capable of discovering, sequencing and genotyping thousands of markers across genomes of interest having prior or no genetic information. In recent years Genotyping-by-sequencing (GBS) has emerged as a promising genomic approach for exploration of plant genetic diversity on a genome-wide scale. There are no studies on assessment of genetic diversity of *Saraca asoca* using genotyping by sequencing (GBS). The GBS approach is suitable for population studies, germplasm characterization, breeding, and trait mapping in diverse organisms.

## 2. Materials and Methods

### 2.1. Study Sites and Plant Sampling

Based on secondary information, databases, and consultations with forest department officers and botanical experts from The University of Trans-Disciplinary Health Sciences and Technology (TDU), wild populations of Saraca asoca were identified in Kollur, Udupi district of Karnataka, India (altitude: 120–49 m amsl, Longitude: 77.43, Latitude: 13.240), as well as in Pune (Longitude: 72.979, Latitude: 19.188) and Bengaluru. The study involved 49 samples of Saraca asoca, with all individual species enumerated and recorded. Fresh leaves, both mature and young, were collected from these identified populations. The leaves were placed in brown covers, sealed in polythene bags with silica adsorbent to remove moisture, and kept on ice until transported to the lab. Once in the lab, the samples were stored in a –80°C freezer.

### 2.2. Isolation of Genomic DNA from Leaf Tissues of *Saraca asoca*

Fresh leaf samples were collected in a sterile bag, leaves were thoroughly washed with distilled water and wiped with ethanol, 100mg of leaf samples were used for genomic DNA extraction. Leaf samples were then finely chopped and ground to fine powder using liquid nitrogen. The lysis buffer was an in-house modified CTAB (Novaes et al. 2009) protocol (2%CTAB, 1.4mM NaCl, 20mM EDTA, 100mM Tris HCl) was used for genomic DNA extraction. To the ground material 1ml of 2% CTAB, 40-60µl of SDS with 60µl of β-Mercaptoethanol was added and were incubated for 1hr at 65°C. The mixture was treated with 24:1 chloroform:isoamyl alcohol, and was centrifuged at 12,000rpm for 15 min, following which the tubes were subjected to ice-cold ethanol and incubated at –20°C for 1hr and was centrifuged at 10,000rpm for 6 min, further the supernatant was discarded, while the pellet was washed with 70% ethanol, following which the pellet was air dried and the DNA pellet was dissolved in 40µl of TE buffer. The DNA sample was quantitated using Nanodrop & Qubit dsDNA BR assay (ThermoScientific, USA). The DNA integrity was checked using a 0.8% Agarose TAE gel.

### 2.3. Library Preparation and Sequencing

Genome sequencing was carried out by Illumina Hiseq2500 Next Generation Sequencing platform. Genomic DNA paired end library preparation was performed through enzymatic reactions, The DNA was enriched with NEBNext Ultra II Q5 master mix, Further the amplified products were cleaned up by using Ampure beads and the final DNA library was eluted in 15 uLs of 0.1X TE buffer. The library concentration was determined by Qubit.3 Fluorometer by Qubit DNA HSassay kit and was quantified using Quant-iTTM DNA HS Assay Kit. Two sequencing paired-end libraries were generated, with 2×150 and 2×100 sequencing chemistry using Illumina HiSeq2500.

For the GBS method, the library preparation was carried out using standard buckler protocol (Elshire et al. 2011). The restriction enzyme used was Pst1 enzyme for digestion, which is a 6-base cutter. After the restriction of digestion, the adapters were ligated to the digested DNA. The samples were further pooled, cleaned and then PCR amplified. The final libraries were validated using the Agilent DNA chip on the Agilent 2100 Bioanalyzer system. Further, the library was quantified using QUBIT dSDNA HS Kit. Samples (49) were digested using the restriction enzyme PstI and sequenced using the Illumina NextSeq 500 with 75 bp single-end reads.

### 2.4. Whole Genome Sequence Analysis

#### 2.4.1. Data QC and *In Silico* Genome Size Estimation

The raw read quality check was carried out with FASTP-v0.20.1 (Chen at al. 2018), low quality reads were trimmed based on phred quality score (phred score >20). The filtered reads were subjected to Insilico genome size prediction through jellyfish v2.2.1 (Marçais and Kingsford 2011) with observed genome peak position at Kmer24. The genome size of *Saraca asoca* was estimated by the formula, genome size = K-mer number/peak depth. The automated estimation of the best K-mer for carrying out genome assembly was performed using KmerGenie v1.7051 (Chikhi and Mahadev 2014).

#### 2.4.2. Genome Assembly

Genome assembly was carried out using both the libraries through SPAdes-V3.11.1 (Bankevich et al. 2012) assembler. Assembly was carried out for a varied range of K-mer size ranging from 21-111. The contigs generated through genome assembly were scaffolded by SSPACE v2.1.1 (Boetzer et al. 2011) and closed the gaps generated through the scaffolding process with GapCloser v1.2.1 (Xu et al. 2020). Genome assembly quality analysis was assessed using QUAST-V5.0.2 (Gurevich et al. 2013), while the genome completeness was assessed by BUSCO (Benchmark Universal Single Copy Orthologs)-v5.2.2 (Sima et al. 2015) through Embryophyta_odb10 lineage datasets.

#### 2.4.3. Repeat identification, Microsatellite SSR Analysis & Primer designing

The assembled genome of *Saraca asoca* was analyzed for the identification of repeat elements using Repeatscout v1.0.5 (Price et al. 2005). The repeats were further masked in the assembled genome through Repeat Masker v4.1.2 (Tarailo-Graovac and Chen 2009). In addition Simple Sequence Repeats (SSRs) or microsatellites survey was carried out using MISAv-1.0.0 (Thiel et al. 2003). The Primer3 (Untergasser et al. 2012) program was used to design PCR primers, in-silico designed primers were subjected to BLAST search for assessing the amplification rate of the primers among closely related plant species. In-vitro validation of the designed primers was performed. The oligonucleotide primer sequence was purified through HPSF inorder to enrich a full length product. The genomic sequences of PCR primers used for validation is given in the (**Table 1**). The amplification of SSR primer was performed in a 25µl of reaction mixture containing 0.4mM dNTP, 1x Taq BufferB, 1.5mM MgCl_2_, 8pmol each of forward and reverse primer, 2 µl of template DNA of *Saraca asoca* and 1.8U Taq DNA Polymerase. The PCR amplification was carried out for 35cycles (3 min at 95 of initial denaturation, 30 s at ℃ of initial denaturation, 30 s at 50.9℃-57.3℃ gradient temperatures for 50.9℃ of initial denaturation, 30 s at 50.9℃-57.3℃ gradient temperatures for-57.3℃ of initial denaturation, 30 s at 50.9℃-57.3℃ gradient temperatures for gradient temperatures for annealing, 1 min 72℃ of initial denaturation, 30 s at 50.9℃-57.3℃ gradient temperatures for extension process) in a 96-well thermocycler. The reaction mixture was further loaded onto 2.5% agarose gel with standard 100bp DNA ladder. The product size of the SSR motif was 275bp, The resultant amplification on gel showed band at 275 bp with respect to 100 bp ladder and presence of shorter bands below 100 bp size.

**Table 1:**
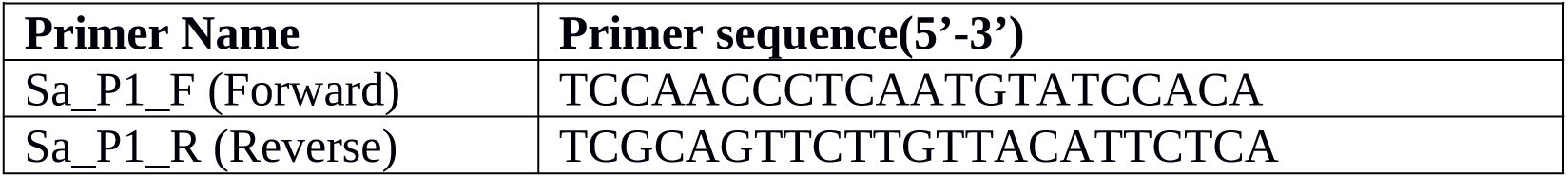
Sequences of SSR primer.

#### 2.4.4. Gene Prediction and Annotation

The gene-structure identification was based on an integrative method with the combination of *ab initio*, homology-based prediction was used in order to predict the gene models for the repeat-masked sequences of *Saraca asoca.* Braker v2.1.6 (Brůna et al. 2021) was used for predicting the genes which uses Gene Mark-EP+ (Brůna et al. 2020) and AUGUSTUS (Stanke et al. 2008). The Braker2 tool uses genomic and protein sequence data for creating the complete gene structure annotations automatically. The mapped *Embryophyta* datasets were used as evidence for gene prediction. Genome annotation and protein-coding genes identification were carried out through the repeat masked genome assembly of *Saraca asoca.* The predicted gene models were aligned with the Uniprot (The UniProt Consortium 2023) and Non-Redundant Protein Sequence Database (NR) (Pruitt et al. 2005) using BLASTX (Camacho et al. 2009). The InterProScan v5.37-76.0 (Jones et al. 2014) was used to identify gene ontology terms for predicted coding genes and visualization through Blast2Go (Conesa et al. 2005). The pathway analysis was carried out through Kyoto Encyclopaedia of Genes and Genomes (KEGG) Automatic Annotation Server (KASS) (Moriya et al. 2007). Further the INFERNAL v1.1.2 (Nawrocki and Eddy 2013) was used for classification of the non-coding RNAs found in *Saraca asoca* genome using the Rfam database (Griffiths-Jones et al. 2003).

#### 2.4.5. Orthology Detection

Some of the species belonging to *Fabaceae* family such as *Phaseolus vulgaris, Vigna radiata, Trifolium pratense, Lupinus angustifolius* were used along with *Saraca asoca* using Orthovenn2 (Xu et al. 2019) to identify orthologous gene clusters among the selected species. Inorder to study the orthologous relationship of *Saraca asoca* across different plant species was carried through Proteinortho (Lechner et al. 2011) with 13 different species: *Bouhinia variegata, Arachis hypogaea, Lupinus angustifolius, Trifolium pratense, Cicer arietinum, Abrus precatorius, Mucuna pruriens, Cajanus cajan, Glycine max, Phaseolus vulgaris, Vigna unguiculata, Vigna umbellata, Vigna radiata.* The proteome sets of all the selected species were obtained from Uniprot. The multiple sequence alignment was performed through MAFFT v7.0 (Multiple Alignment using Fast Fourier Transform; Katoh et al. 2002) and the results were subjected to GBLOCKS v0.91b (Castresana 2000) program for processing the MAFFT alignment. The processed alignment was used for phylogenetic analyses of large datasets under maximum likelihood through RAxML v1.0.0 (Stamatakis 2014) from all the 14 different species.

#### 2.4.6. GBS Data Analysis

The reads underwent a 3’-end trimming process utilizing Trimmomatic v. 0.3 (Bolger et al. 2014). This trimming aimed to attain an average Phred quality score of ≥20 across a ten-base window. Additionally, reads with a final length of less than 20 base pairs were excluded. For GBS analysis the assembled genome of *Saraca asoca* was used as a reference genome. BWA-MEM v0.7.17 (Li and Durbin 2009) was used for mapping of quality trimmed reads against reference genome. Samtools (Danecek et al. 2021) was used to preprocess the aligned reads. Variant discovery and genotype calling were performed using Freebayes v. 1.2.0 (Garrison and Marth 2012). The SNPs were filtered based on the criteria such as a minimal read count > 8, a minimum allele frequency (MAF) ≥ 0.05, and proportion of samples with designated genotype ≥ 66%. Genetic diversity analyses were conducted employing biallelic SNPs possessing complete genotypic information across the entirety of samples, without any instances of missing data.

To optimize data analysis and achieve a more profound comprehension of the intricacies underlying genetic diversity patterns, the accessions were grouped based on their respective geographic locales **(Supplement-1; Table-1).** The genetic diversity within the 49 accessions was evaluated through the utilization of diverse statistical methodologies. Further downstream analysis such as analysis of heterozygosity, diversity analysis, PCA analysis, population analysis was carried out using SNiPlay3 (Dereeper et al. 2015)

## 3. Results

### 3.1. Genome Sequencing and De novo Genome Assembly of *Saraca asoca*

Paired end reads of two libraries with 10x and 100x coverage, with sequencing chemistry (2×150 bp) were generated using Illumina Hiseq 2500. Sequencing yielded a combined total of 475.98 million reads with 100x libraries and 120 million reads with 10x libraries. After filtering a total of 460M reads and 118.82M reads were passed through the quality check for 100x and 10x libraries respectively **(Supplement-2; Table 1)**. Genome size estimation was conducted by considering reads with a quality score greater than 20. This analysis revealed a haploid genome length of 1,602,089,559 bases, a genome repeat length of 746,525,378 bases, and a genome unique length of 855,564,181 bases **(Supplement-2; Table 2)**. The best k-mer size for assembly was predicted using Kmergenie which predicted the assembly size of 2, 044,244,623 bp (∼ 2Gb). The genome assembly with k-mer size 85 resulted in a genome size of 2,012,082,659 bp. The contigs from the assembly were scaffolded which obtained a maximum contig length of 86,394 bp with N50 of 5,066 bp. The gaps in the Scaffolding were closed through Gapcloser resulting in the gapclosed genome size of 1,651,830,046 bp **(Table 2)**. Genome completeness was assessed through BUSCO resulting in 1181 (73.2%) complete BUSCOs (C), 1,110 (68.8%) Complete and Single-copy BUSCOs (S), 71 (4.4%) Complete and Duplicated BUSCOs (D), 268 (16.6%) Fragmented BUSCOs (F) and 165 (10.2%) Missing BUSCOs (M) in a total of 1,614 BUSCO groups searched **(Supplement-2; Table 3)**.

**Table 2:** Genome assembly statistics of *Saraca asoca*.

| Assembly parameters | Assembly statistics |
| --- | --- |
| Total high quality reads | 1,651,830,046 |
| K-mer (bases) | 85 |
| **Assembled genome size (Gb)** | 1.6 |
| **Total number of contigs** | 767,695 |
| **N50 (bases)** | 5,066 |
| **Maximum contig length (bases)** | 86,394 |
| **Minimum contig length (bases)** | 460 |
| **% of bases in contigs ≥ 1,000 bases** | 82.2% |
| **Total repeat size (Mb)** | 746.53 |

**Table 3:** Genome annotation statistics of *Saraca asoca*.

|  |  |
| --- | --- |
| **Number of genes** | 63,913 |
| **Number of cds** | 63,913 |
| **Number of exons** | 198,458 |
| **Number of introns** | 151,161 |
| **Number of start codons** | 46,634 |
| **Number of stop codons** | 50,012 |
| **Total gene length (bases)** | 101,560,147 |
| **Longest gene (bases)** | 43,659 |
| **Longest exon (bases)** | 7,962 |
| **Longest intron (bases)** | 11,933 |

### 3.2. Genome Annotation

The genome annotation was carried out using Repeat masked genome assembly. An integrated approach, combining *ab initio* and homology-based methods, was employed for gene prediction to identify protein-coding genes. Total of 63,913 protein coding genes were identified, with 1,98,458 exons. Average gene length of 1,589 bp, with an average of 3.1 exons per cds, covering average exon length to be 268 bp **(Table 3)**. The functional annotation of *Saraca asoca* was performed by aligning predicted gene models against different databases such as uniprot, nr, KEGG and Interproscan. A total of 62,038 genes were functionally annotated by the nr database, 62,038 genes were functionally annotated from the uniprot database. The gene ontology analysis resulted in 13,000 genes carrying out various biological processes (BP), around 15,000 genes in various molecular functions (MF), and around 6,000 genes for cellular components (CC) as shown in **(Supplement-3: Figure 1)**. The majority of genes in biological processes are involved in cellular processes, metabolic processes, biological regulation, response to stimulus, *etc*. While genes coding for molecular functions are involved in binding, catalytic activity, ATP dependent activities, structural molecular activity and so on, the genes carrying out cellular components have shown the enriched function of cellular anatomical activity, protein containing complex *etc*.

*Saraca asoca* gene models were searched against the Pfam database to identify protein families and domains. Of 63,913 protein coding genes, 32,453 gene models have been identified with at least one Pfam domain. Total 4,332 Pfam domains were identified. The pfam domain Pkinase (PF00069.24) is the major domain and is identified in 1,527 genes which is followed by Pkinase_Tyr ( PF07714.16), LRR_8 (PF13855.5), PPR (PF01535.19), LRR_4 ( PF12799.6), etc were identified in 1,492, 725, 688, 687 gene models, respectively **(Supplement-2; Table-5)**. The gene containing the highest pfam domain was further analyzed. This protein associated with the protein family “Midasin-related,” exhibits a diverse array of domains. Among these are the AAA domain (dynein-related subfamily; PF07728) and the AAA_5 domain (SM00382). Additionally, the protein is characterized by the presence of the Midasin AAA lid domain (PF17865 and PF17867) These domains suggest involvement in processes such as ATP-binding and nucleoside triphosphate hydrolase activity, as well as potential interactions with sigma-54. Moreover, the protein displays sequence motifs consistent with the AAA family (cd00009) **(Supplement-3: Figure 2)**.

**Figure 2:**
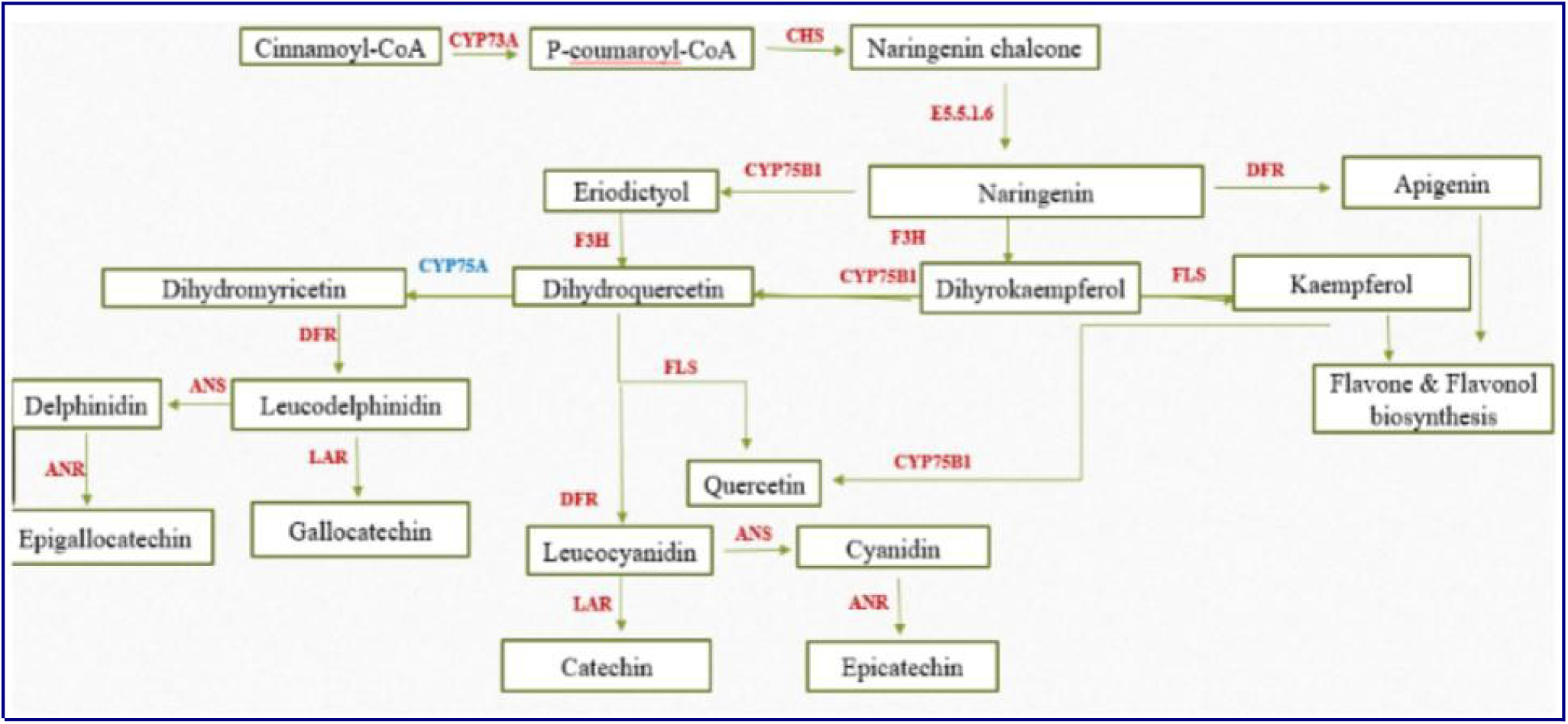
Biosynthesis pathway map of different flavonoids such as catechin, epicatechin, quercetin and epigallocatechin from *Saraca asoca* genome.

### 3.3. Pathway Analysis

Reports indicated that preliminary phytochemical analysis of various extracts (bark, flower, and leaves) from *Saraca asoca* was conducted, revealing the presence of Alkaloids, Flavonoids, Glycosides, Saponins, Phenols, Steroids, Tannins, and Triterpenoids [6]. The bark of *Saraca asoca* is known to have stimulating effect on endometrium, ovarian tissues useful in treating menorrhagia, leukorrhea and other gastrointestinal conditions. Various parts of *Saraca asoca* exhibit a diverse array of reported metabolites, including the presence of catechin, epicatechin, and epigallocatechin (Shirolkar et al. 2013). Initially, Cytochrome P450 73A (CYP73A) converts cinnamoyl-CoA to P-coumaroyl-CoA, utilized by Chalcone synthase (CHS) to produce naringenin chalcone, a crucial intermediate. Subsequently, the enzyme with the code E5.5.1.6 (E5.5.1.6) and Dihydroflavonol 4-reductase (DFR) play vital roles in further modifying the intermediates, leading to the production of dihydroflavonols. These dihydroflavonols undergo hydroxylation reactions mediated by enzymes like Cytochrome P450 7581 (CYP7581), Cytochrome P450 75A (CYP75A), and Cytochrome P450 75B1 (CYP75B1). Flavanone 3-hydroxylase (F3H) catalyzes the conversion of flavanones to dihydroflavonols, a pivotal step in the pathway. Further downstream, Flavonol synthase (FLS) facilitates the conversion of dihydroflavonols to flavonols, including quercetin, contributing to the diverse array of flavonoids synthesized. As the pathway progresses, Anthocyanidin synthase (ANS) transforms colorless leucocyanidin and leucodelphinidin into colored anthocyanidins, while Anthocyanidin reductase (ANR) and Leucoanthocyanidin reductase (LAR) reduce anthocyanidins to flavan-3-ols like catechin and epicatechin **(Figure 2)**.

### 3.4. Orthology Detection

Orthology detection was carried out through Orthovenn2. The predicted protein coding genes of *Saraca asoca* were analyzed through Orthovenn2, along with closely related species to that of *Saraca asoca* for orthologous cluster analysis. The occurrence table depicted the pattern of occurrence between shared orthologous groups among *Saraca asoca, Phaseolus vulgaris, Vigna radiata, Trifolium pratense, Lupinus angustifolius.* The cluster count shows the number of clusters shared among different species, with 4,975 clusters shared among all the five species including 32808 proteins **(Supplement-3: Figure 3)**. The Orthology detection resulted in 63,913 protein members in the shared clusters of *Saraca asoca,* with the highest number of clusters to be 19,971 for *Saraca asoca.* A total of 29,668 singletons specific to the species of *Saraca asoca* were identified. The *Saraca asoca* shares more clusters with *Phaseolus vulgaris*. The *Saraca* genome consists of 4812 paralog clusters and 28000 singletons **(Supplement-3: Figure 4)**. The phylogenetic analysis was carried out among various *Fabaceae* family members, which resulted in phylogenetic association representing evolutionary close relation to *Bauhinia variegata & Arachis hypogaea* with its evolutionary divergence value represented on the branches of phylogenetic representation; The tree also shows evolutionary relationship among all the closely related species **(Figure 3)**.

**Figure 3:**
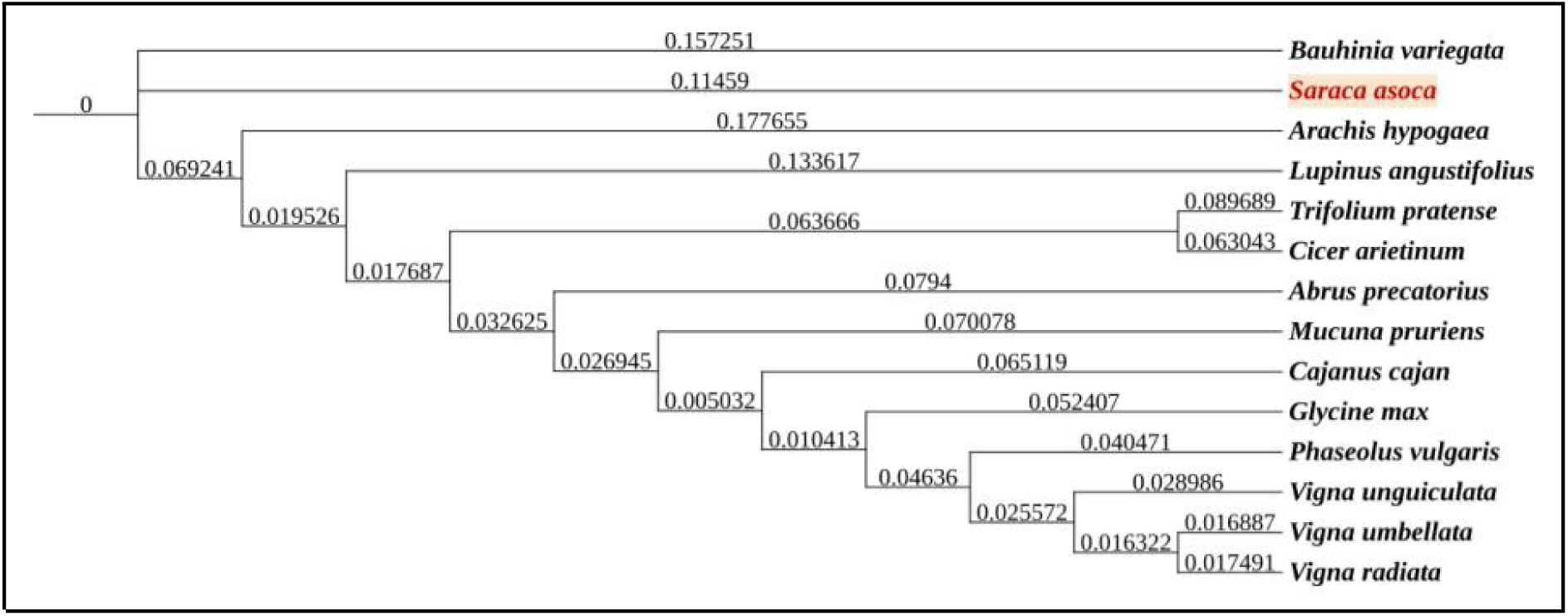
Phylogenetic tree constructed from of *Saraca asoca* proteome.

**Figure 4:**
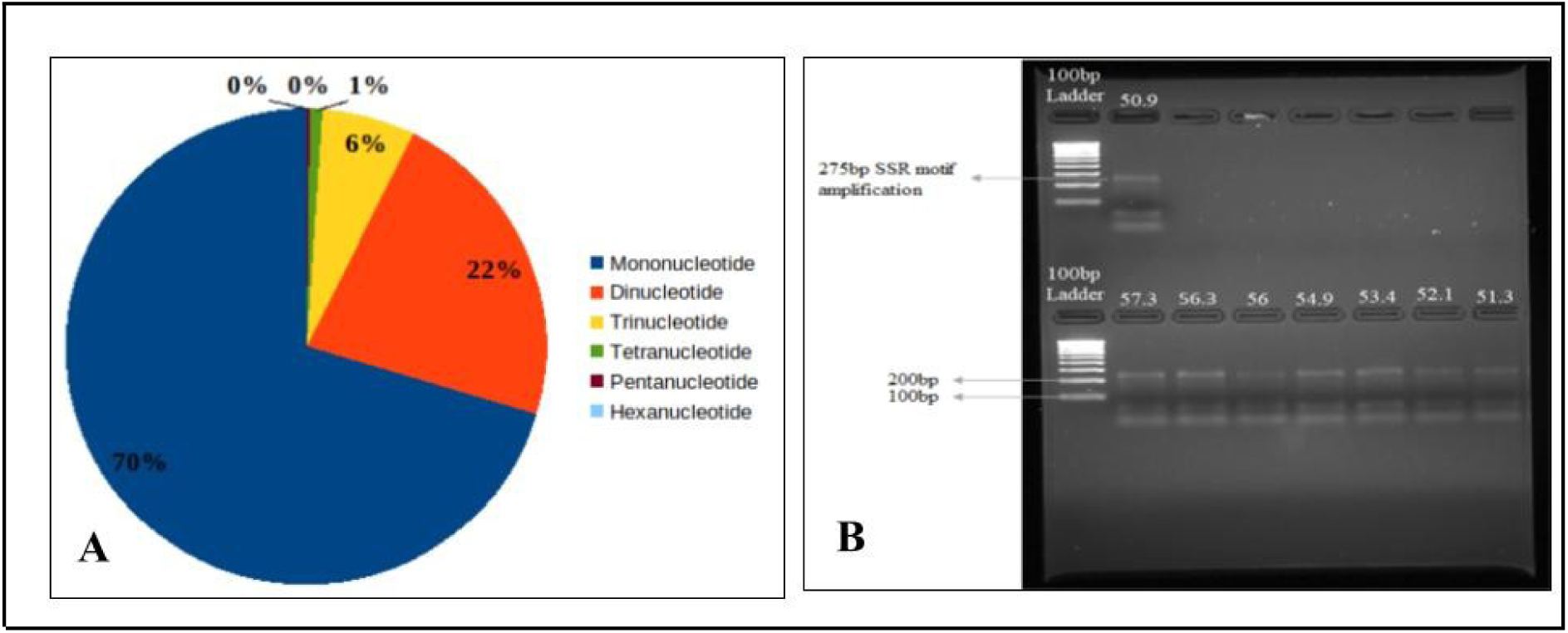
Analysis of Simple Sequence Repeats (SSRs) in the *Saraca asoca* genome. A – Distribution of SSR types based on repeat unit length; B – PCR amplification of a 275 bp SSR motif using specific primers. The gel electrophoresis image shows amplified SSR products from different Saraca asoca samples, with a 100 bp DNA ladder used for size comparison.

### 3.5. SSR Analysis

A Simple Sequence Repeat (SSR) survey was carried out for the *Saraca asoca* genome assembly, identifying a total of 5,84,615 SSRs across 2,36,123 SSR-containing sequences. The analysis encompassed 7,67,671 sequences, with a total examined size of 1,651,794,440 base pairs. A significant proportion of the sequences (2,36,123) contain SSRs, with 1,10,191 sequences harboring more than one SSR. Additionally, 65,639 SSRs were identified in compound formation, indicating a high level of complexity **(Table 4)**. The table also shows that mononucleotide repeats were the most prevalent, constituting 70.47% of the total SSRs, followed by dinucleotide (22.15%) and trinucleotide (6.27%) repeats. Smaller proportions of tetra, penta, and hexanucleotide repeats were also observed **(Figure 4A)**.

**Table 4:** Microsatellite (SSR) analysis.

| Results of Microsatellite(SSR) search | Statistics |
| --- | --- |
| Total number of sequences examined | 7,67,671 |
| Total size of examined sequences (bp) | 1,651,794,440 |
| Total number of identified SSRs | 5,84,615 |
| Number of SSR containing sequences | 2,36,123 |
| Number of sequences containing more than 1 SSR | 1,10,191 |
| Number of SSRs present in compound formation | 65,639 |

The analysis of dinucleotide SSR motifs in the *Saraca asoca* genome reveals that the most prevalent motifs are “AT” (36.53%) and “TA” (23.45%), collectively accounting for nearly 60% of all dinucleotide SSRs. This dominance suggests a preference for AT-rich regions in the genome. In contrast, motifs like “CG” and “GC” are the least abundant, each contributing less than 0.1%, which is consistent with the typically low occurrence of GC-rich SSRs in plant genomes **(Supplement-2: Table 4).** The analysis of trinucleotide SSR motifs in the *Saraca asoca* genome indicates that “AAT” (13.96%) is the most abundant motif, followed by “TTA” (10.57%) and “ATT” (10.57%). These motifs collectively constitute a significant portion of the trinucleotide repeats, highlighting a preference for A/T-rich sequences. In contrast, motifs such as “CGA” (0.016%) and “ACG” (0.027%) are among the least represented, consistent with the lower frequency of GC-rich trinucleotide repeats in plant genomes **(Supplement-2: Table 5)**.

### 3.6. Primer Designing

The automated SSR marker development was carried out through the Primer3 program to design PCR primers for the obtained SSR markers of *Saraca asoca.* The primer3 resulted in three sets of primer pairs for each identified SSR motifs in the complete genomic sequences of *Saraca asoca.* The amplification rate of the primers among closely related plant species was assessed through BLAST search. Thus the in-silico amplifying primers with suitable melting temperature, product size, and best blast hits were selected for In-vitro validation. The In-vitro PCR amplified results were documented as shown below in (**Figure 4B**). The product size of the SSR motif was 275 bp, The resultant amplification on gel showed band at 275 bp with respect to 100 bp ladder and presence of shorter bands below 100 bp size.

### 3.7. Genetic Diversity Analysis of *Saraca asoca*

#### 3.7.1. Identification of SNPs

The quantification of the genomic DNA sample was done using Nanodrop and was checked for the quality using 260/280 ratios and 260/230 ratios **(Supplement-1; Table 1)**. For the 49 samples, the total number of reads ranged from 2.7 million to 23.1 million with an average of 9 million reads per sample. The total number of bases sequenced in these 49 samples was 63.7 Gb with an average of 1.3 Gb per sample. The reads were filtered for low quality reads. After filtering, 479 million reads were obtained for all 49 samples ranging from 2.7 million to 23.1 million reads with an average of 9.3 million reads per sample. Total 62.9 Gb data was obtained for all 49 samples after filtering ranging from 377.9 Mb to 3.2 Gb with an average of 1.3 Gb per sample **(Sapplement-1; Table 3)**.

The alignment of reads obtained from 49 *Saraca asoca* genotypes led to the identification of 560,044 single nucleotide polymorphisms (SNPs). Further SNPs were filtered based on MAF above 5% and without missing data resulted in 9414 SNPs. These SNPs were further filtered for biallelic SNPs which resulted in 8738 SNPs and these SNPs were used for genetic diversity analysis across 50 *Saraca asoca* genotypes. A total of 8738 SNPs were distributed among 938 scaffolds. Notably, approximately 4.9% of these scaffolds exclusively contained a single SNP, despite the fact that the number of SNPs per scaffold varied, ranging from 1 to as many as 76. The mean and median of the MAF of the SNPs were 0.53 and 0.32, respectively, The polymorphism information content (PIC) of the SNPs varied from 0 to 0.5 with a mean and third quartile of 0.12 and 0.28, respectively **(Supplement-3: Figure 5)**.

#### 3.7.2. Genetic Diversity of the *Saraca asoca* Genotypes

The analysis reveals a distinct clustering pattern of *Saraca asoca* genotypes based on their populations. Genotypes from the same population categories, namely Pop-I, Pop-II, Pop-III, and Pop-IV, tend to exhibit similar patterns in the principal component space. This clustering suggests a significant genetic coherence within populations, reflecting shared genetic characteristics or adaptations. The principal components provide a quantitative representation of the observed variations, emphasizing the genetic diversity and population structure of *Saraca asoca* genotypes.

Genetic diversity within and between populations by analyzing heterozygous sites across *Saraca asoca* individuals sampled from three distinct populations (Pop-I, Pop-II, and Pop-III) based on geographical location. Within Pop-I, the maximum observed percentage of heterozygous sites is 43.54%, while the minimum is 20.19%. Similarly, in Pop-II, the range spans from 20.19% to 27.58%. Comparatively, Pop-III exhibits a narrower range, with values ranging from 20.19% to 25.86% suggesting relatively lower genetic diversity within these populations compared to Pop-I **(Supplement-3: Figure 6)**. Tajima’s D value was calculated on a 100 k sliding window. The observed Tajima’s D diversity values, calculated for a set of high-confidence SNPs, demonstrated significance and a predominantly positive trend. Among the 911 calculated Tajima’s D values, a notable majority, comprising 742 instances, exhibited positive values, indicating an excess of intermediate-frequency alleles within the population. Conversely, 169 Tajima’s D values were negative, suggesting potential deviations from neutrality in the genetic variation observed at these specific loci. This skew towards positive Tajima’s D values suggests potential selection pressures or demographic events shaping the genetic diversity of the population **(Supplement-3: Figure 7).** The phylogenetic tree illustrates the population structure and genetic diversity within and between the identified populations, denoted as Pop-I, Pop-II, and Pop-III. Pop-I exhibits a structured arrangement, with several subclusters indicating distinct genetic lineages within this population. In contrast, Pop-II displays a more dispersed pattern, suggestive of greater genetic diversity and potential admixture. The branches representing Pop-II show varying lengths, implying differential genetic distances among its constituent samples. Pop-III appears as a distinct cluster, diverging from both Pop-I and Pop-II, indicative of a separate genetic lineage **(Figure 5A)**. To further validate the population structure, cross-entropy analysis highlights K=4 as the optimal number of clusters, with the lowest cross-entropy value observed at this point **(Figure 5B)**. This supports the distinct groupings of genotypes into three primary populations, with potential sub-structuring within Pop-I. Principal Component Analysis (PCA) further corroborates these findings, as the PCA plot clearly separates the three populations along the first two principal components **(Figure 5C)**. Pop-I and Pop-III form tightly clustered groups, reflecting their relatively homogenous genetic makeup, whereas Pop-II is more scattered, indicating greater genetic diversity and possible admixture.

**Figure 5:**
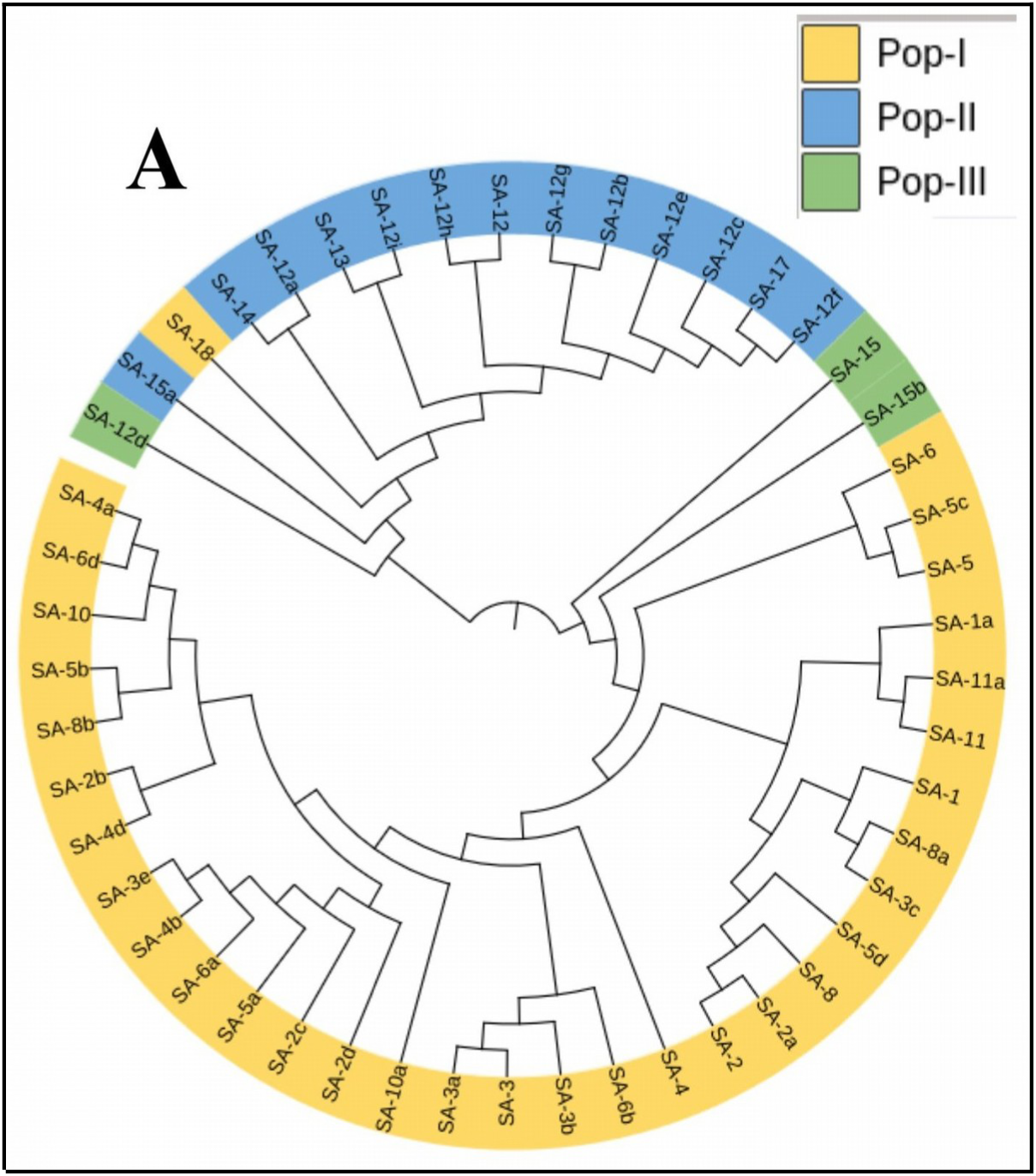
Genetic diversity and population structure in 49 *Saraca asoca* Genotypes. A-Phylogenetic tree showing the clustering of 49 genotypes into three distinct populations (Pop-I, Pop-II, and Pop-III) based on genetic distance. Populations are color-coded: yellow (Pop-I), blue (Pop-II), and green (Pop-III)

**Figure 5:**
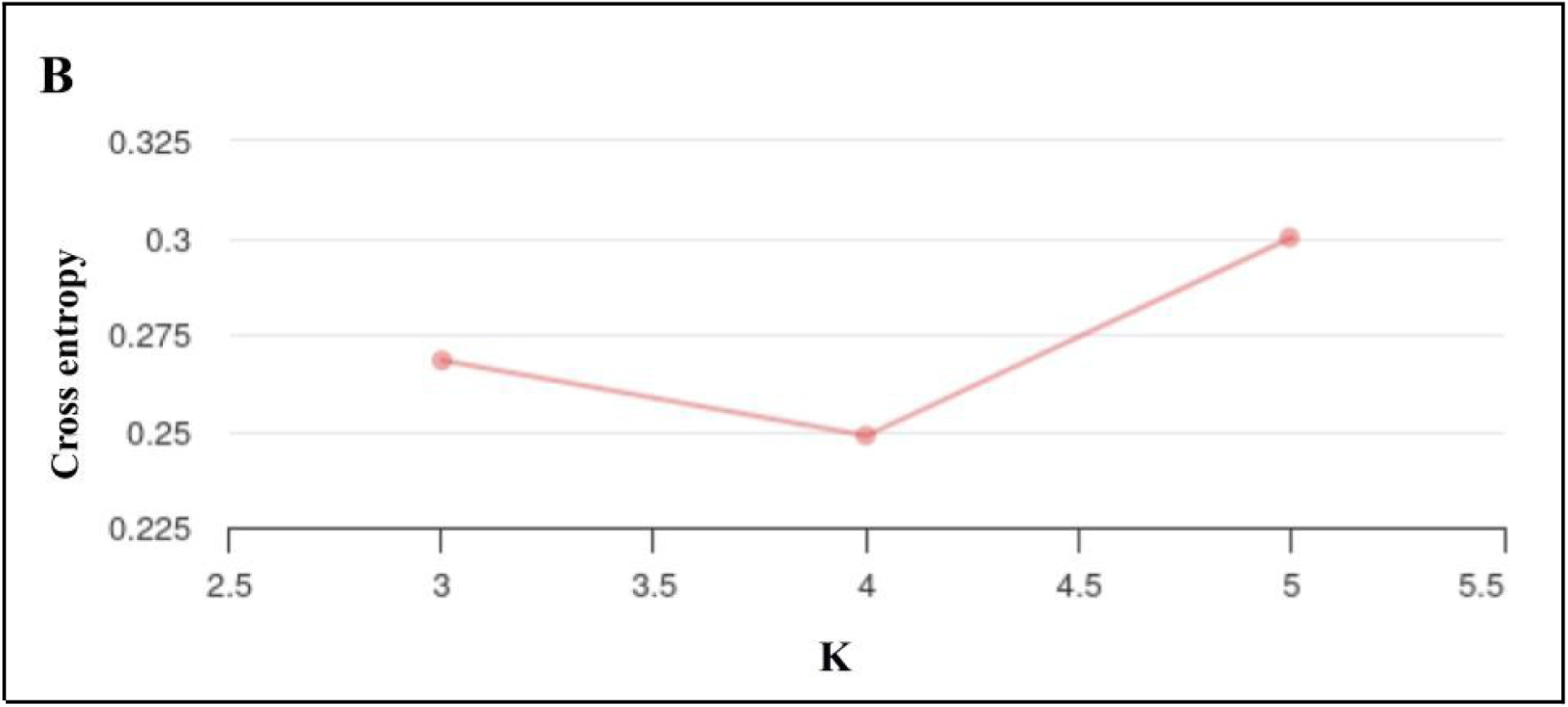
Genetic diversity and population structure in 49 *Saraca asoca* Genotypes. B – Cross-entropy plot showing the optimal value of K for population structure analysis, with the lowest cross-entropy observed at K=4, indicating the best fit for the genetic data

**Figure 5:**
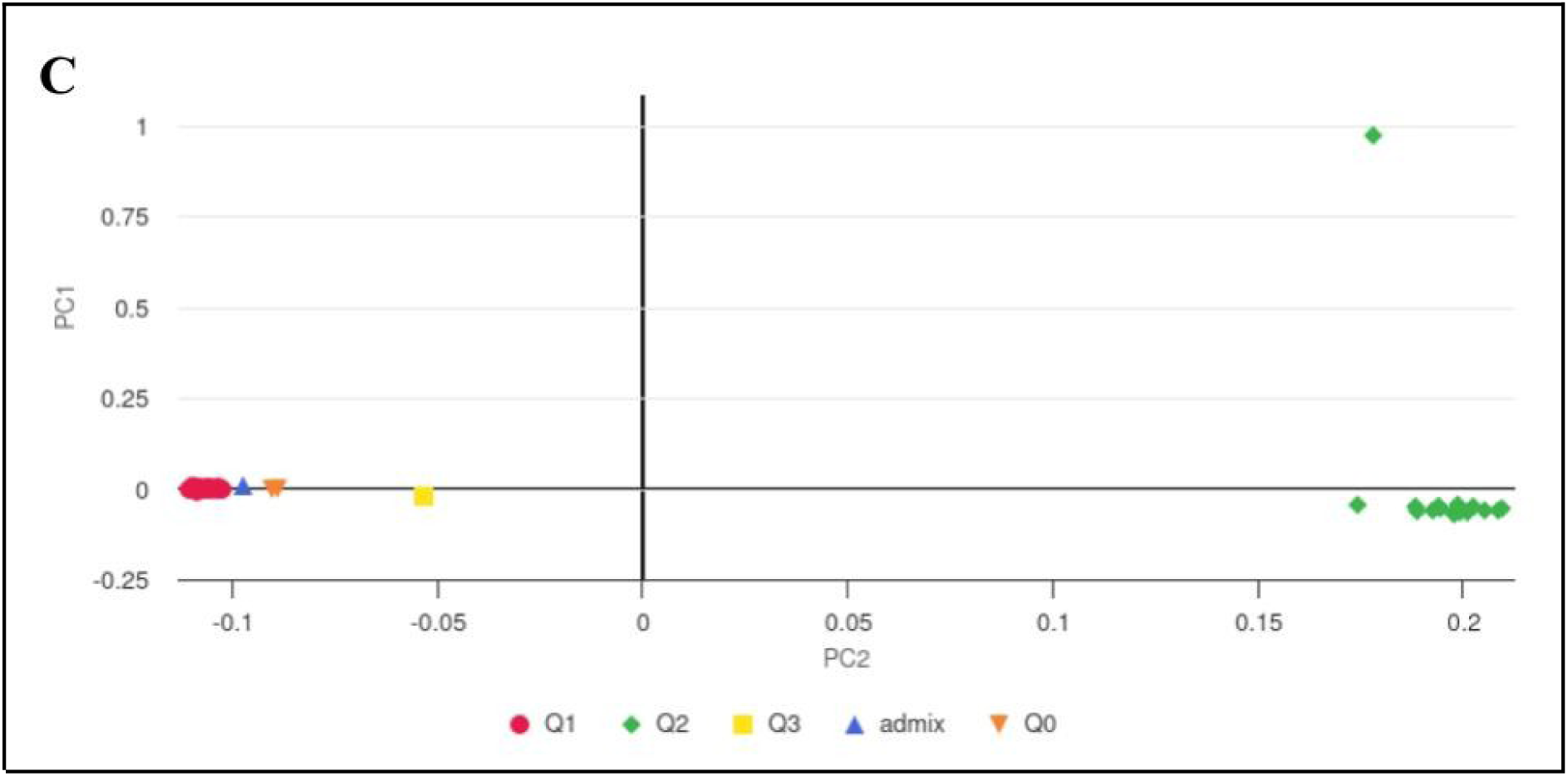
Genetic diversity and population structure in 49 *Saraca asoca* Genotypes. C – Principal Component Analysis (PCA) plot illustrating the genetic variation among the genotypes. Points are color-coded by population, and the clustering corresponds to the groupings observed in the phylogenetic tree. Outliers and admixture are also indicated.

**Figure 6:**
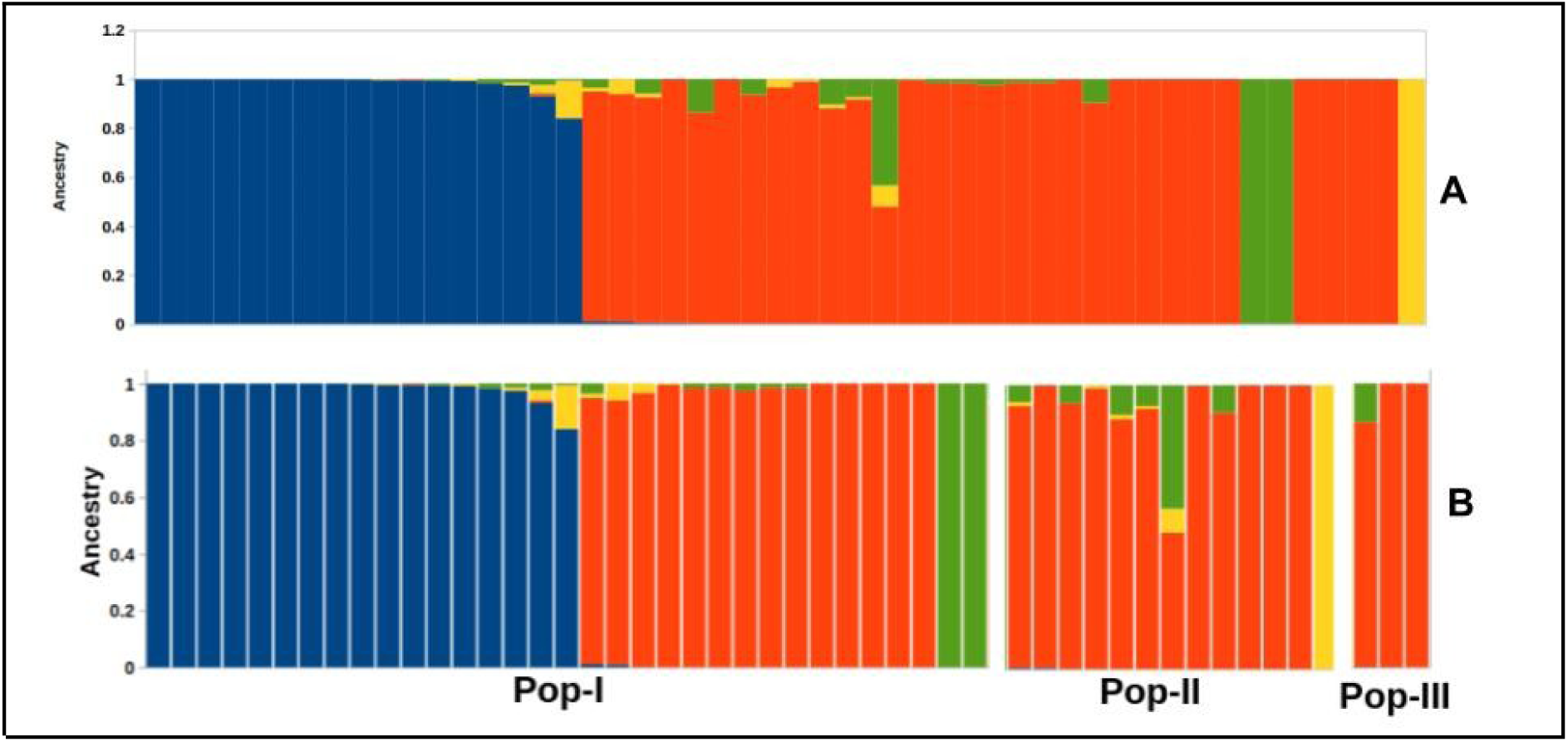
A visual representation of the genetic composition of 49 Saraca asoca samples at K = 4 reveals varying degrees of admixture. (A) Samples were organized based on their membership levels in distinct clusters, while (B) individual accessions were sorted according to their designated populations.

#### 3.7.3. Population Structure Analysis

The population genetic structure analysis, based on 8738 SNPs, revealed three distinct genetic populations within the 49 accessions studied. Pop-I predominantly consists of samples with high membership probabilities in Cluster I, indicating a strong genetic affinity to this ancestral group. In contrast, Pop-II exhibits a predominant association with Cluster II, suggesting distinct genetic characteristics from Pop-I. Pop-III, on the other hand, shows a clear preference for Cluster III, delineating its unique genetic composition. This organization highlights the presence of distinct genetic clusters within the *Saraca asoca* samples, suggesting differentiated population structures corresponding to Pop-I, Pop-II, and Pop-III **(Figure 6A and 6B).** Further, diagnostic SNPs were identified using SNiPlay3, a tool for population genetics analysis. A total of 40 diagnostic SNP markers were identified across 36 scaffolds in the Saraca asoca genome. Gst values, a measure of genetic differentiation, were calculated for each SNP marker to determine their population specificity **(Supplement 1; Table 4).** Among these SNPs, 12 were specific to Pop-I, 24 to Pop-II, and 4 to Pop-III. The Gst values ranged widely, indicating varying levels of differentiation across the populations. Certain SNPs exhibited high Gst values in specific populations, such as 0.942 for Pop-I and 0.883 for Pop-II, underscoring their potential utility in distinguishing these populations.

## 4. Discussion

The major threat to the world is the loss of wild plant biodiversity that is drastically increasing over time due to various reasons. Species reduction has been observed to be as 1000 times higher than background due to human actions. Habitat loss and fragmentation, overexploitation, pollution, invasion of alien species, global climatic changes (IUCN (2011) IUCN red list of threatened species. Version 2011.1, Smitha et al. 2016) and disruption of community structures has been found to have major threats to ecosystems (Shukla 2017). This plant has been harvested unsustainably for medicinal use and therefore it has attained the status of ‘Endangered’ (IUCN (2011) IUCN red list of threatened species. Version 2011.1). Limited efforts have been taken to conserve in insitu and exsitu. Therefore there is a need for method development for in situ conservation of *Saraca asoca* genetic resource. This study was taken to assess the genetic diversity between *Saraca asoca* samples collected from MPCA, Kolluru for identification of most diverse population, which can be conserved, and germplasm of the same may be used for propagation and cultivation

*Saraca asoca* is a medicinally, environmentally essential tropical Indian tree species. *Saraca asoca* is reported to contain glycosides, flavonoids, tannins and saponins attributing to its potential for treating various gynecological issues with spasmogenic, oxytocic, uterotonic, anti-bacterial, anti-tumour, anti-progestational, anti-estrogenic activity against menorrhagia and anti-cancer properties (Pradhan et al. 2009). The bark of *Saraca asoca* is one of the highly traded herbs in the ayurvedic and herbal industry being an essential ingredient in almost all the ayurvedic formulations for treating gynecological problems (Shukla 2017). However molecular, genetic and genomic research are not well studied to understand genes, genomic analysis of *Saraca asoca.* Recently, studies on SSRs in the chloroplast genome of *Saraca asoca* were generated (Ali et al. 2021). However these studies have generated SSRs from the chloroplast genome. Our research intends to develop the whole genome assembly, genome annotation, pathway analysis of certain essential metabolites with whole genome SSR marker development study of *Saraca asoca* which can aid in genetic diversity studies for conservation of the vulnerable species.

We sequenced *Saraca asoca* genome collected from The University of Transdisciplinary Health Sciences and Technology (TDU), through next generation sequencing platform from Illumina Hiseq-2500 at Bangalore Genomics Centre. We could assemble 76% (1.6 Gb) of genome based on estimated genome size for *Saraca asoca.* More than 73% of conserved genes mapped to the de novo assembled genome of *Saraca asoca.* Our analysis predicted the total repeat size to be 746 Mb from the *Saraca asoca* genome, with identification of molecular markers such as SSRs from the assembled whole genome of *Saraca asoca*.

Our study added whole genome assembly, genome annotation, molecular marker study aiding in detailed metabolic pathway analysis from *Saraca asoca* in future. We identified genes involved in the biosynthesis of secondary metabolites Catechin, Epicatechin, from the flavonoid synthesis pathway from KEGG. predicted the total repeat size to be 746 Mb in *Saraca asoca* genome. The identified genes from this study can enhance research on essential metabolite biochemical pathways. The genetic diversity data from our study can aid in the conservation of vulnerable tree species *Saraca asoca*

Shirolkar et al. (2013) reported that catechin and epicatechin were upregulated in the bark and leaves of *Saraca asoca*, while epigallocatechin was found to be in higher concentration in the bark water extract in comparison to other parts of *Saraca asoca* with considerably lower antimicrobial properties, thus catechin and epicatechin are well known flavonoids attributing to the antioxidant activity, leading to the asymptomatic treatment of various vascular, respiratory and gastrointestinal disorders (Shirolkar et al. 2013).

The genetic diversity analysis of *Saraca asoca* conducted in this study sheds light on the intricate genomic landscape of this medicinal plant species. Through the identification of single nucleotide polymorphisms (SNPs), the study uncovered a total of 8738 biallelic SNPs distributed across 938 scaffolds, indicative of considerable genetic variation within the sampled population. Notably, the majority of scaffolds contained multiple SNPs, with a small proportion exhibiting exclusive single SNP occurrences. The observed mean minor allele frequency (MAF) of 0.53 underscores the presence of common genetic variants within the population, while the polymorphism information content (PIC) values provide insights into the informativeness of the SNPs for genetic diversity assessment. The distinct clustering patterns observed among *Saraca asoca* genotypes from different populations highlight the population structure and genetic coherence within each group. Additionally, Tajima’s D analysis revealed predominantly positive values, suggesting potential selection pressures or demographic events shaping the genetic diversity of the population. The identification of diagnostic SNP markers further enriches our understanding of population genetics and offers valuable insights into population differentiation within *Saraca asoca*.

These results underscore the significance of genetic diversity analysis in elucidating the population structure and evolutionary dynamics of *Saraca asoca* (Hamrick and Godt 1996). The observed clustering patterns and population differentiations highlight the genetic coherence within populations and provide valuable information for conservation and breeding programs aimed at preserving and enhancing the genetic diversity of this medicinally important plant species (Pritchard et al. 2000, Evanno et al. 2005). Furthermore, the identification of diagnostic SNP markers offers potential applications in marker-assisted breeding and population genetic studies, facilitating targeted selection of desirable traits and genetic improvement strategies (Rafalski 2002, Mammadov et al. 2012).

However, it is essential to acknowledge the limitations of this study, including the relatively small sample size and the potential influence of sampling bias on the observed genetic patterns. Further studies incorporating larger sample sizes and additional genomic markers could provide a more comprehensive understanding of Saraca asoca’s genetic diversity and population structure. In conclusion, these findings contribute to our knowledge of *Saraca asoca*’s genetic landscape and lay the groundwork for future research aimed at leveraging genetic resources for conservation and sustainable utilization of this valuable medicinal plant species.

## 5. Conclusion

The sequencing of *Saraca asoca* genome from the Illumina Hiseq-2500 resulted in genome assembly of size 1.6Gb. More than 73% of conserved genes mapped to the de novo assembled genome of *Saraca asoca.* We annotated the *Saraca asoca* genome to the best of our knowledge through available bioinformatics tools. Adding on the SSR marker development from the assembled whole genome of *Saraca asoca.* Thus the current research can lead to availability of Whole genome sequencing and analysis of *Saraca asoca* to the public domain, SSR survey of *Saraca asoca* genome with biosynthesis pathway analysis of some of the essential metabolites such as Catechin, Epicatechin, attributing to the rich source of medicinal properties of *Saraca asoca*.

## 6. Data Availability

The raw sequence reads used for whole genome assembly are deposited in NCBI SRA database under bioproject PRJNA770455. The whole genome assembly has been deposited at DDBJ/ENA/GenBank under the accession JANHAO000000000

## 7. Acknowledgements

Authors acknowledge Rural India Supporting Trust (RIST) for the financial support

